# Osmotic fragility during *in vitro* erythrocyte cytotoxicity induced by aluminium chloride, lead acetate or mercuric chloride in hyposmolar sucrose media

**DOI:** 10.1101/2022.12.03.515355

**Authors:** Nanacha Afifi Igbokwe, Ikechukwu Onyebuchi Igbokwe

**Affiliations:** Department of Veterinary Physiology and Biochemistry, Faculty of Veterinary Medicine, University of Maiduguri, P. M. B. 1069, Maiduguri, Nigeria; Department of Veterinary Pathology, Faculty of Veterinary Medicine, University of Maiduguri, P. M. B. 1069, Maiduguri, Nigeria

**Author notes:** Corresponding author, Ikechukwu Onyebuchi Igbokwe, Strategic Animal Research Group, Faculty of Veterinary Medicine, University of Maiduguri, PO Box 8000, Maiduguri, Nigeria.

**Keywords:** erythrocyte death, erythrocyte osmotic fragility, inducible osmotic resistance, metallic salts, xenobiotic cytotoxicity

## Abstract

Erythrocyte death by eryptosis or erythronecrosis may induce erythrocyte shrinking or swelling with increase in osmotic resistance or fragility as indication of cytotoxicity. We investigated heterogeneous cytotoxic outcomes during in vitro exposure of goat erythrocytes to aluminium chloride, lead acetate or mercuric chloride using erythrocyte osmotic fragility (EOF) testing. The metallic salt solution (MSS) was added to 4.0 μL of high (100 mosmol/L) and low (250 mosmol/L) hyposmolar sucrose media at 0.3 or 0.4 mosmol/L concentration during testing of the osmotic fragility of 5.0 μL of blood from 10 goats. Hemolysis induced in the media (with and without MSS) was estimated in the supernatant with spectrophotometer at 540 nm. Osmotic stabilization or destabilization was calculated with probability for each test. Inducible osmotic resistance (IOR) was the ratio of mean stabilization to destabilization in both high and low hyposmolar media. Each MSS induced both osmotic resistance (stabilization) and fragility (destabilization) in varied media concentrations, with greater likelihood (P) of stabilization (0.93) or destabilization (0.77) in high or low media hyposmolarity, respectively. The EOF outcomes of the goats diverged within the group. High IOR induced by mercuric chloride (2.90) and low IOR by lead acetate (0.07) and aluminium chloride (0.04) reflected high stabilizing and destabilizing outcomes, respectively. In conclusion, MSS induced dual EOF outcomes (stabilization or destabilization) on the fragility domain, suggesting occurrence of both eryptosis (as stabilization) and erythronecrosis (as destabilization) at low exposure level, whereby biphasic, nonmonotonic or hormetic response to MSS toxic action might exist.

## Introduction

Aluminium (Al), lead (Pb) and mercuric (Hg) compounds are inorganic xenobiotics and metallic toxicants to humans and animals causing erythrocyte damage or death in the bloodstream (Pagano and Faggio 2015). Anemia is an important toxic effect arising from their toxicosis (Flora *et al*., 2012; Wani *et al*., 2015; Igbokwe *et al*., 2019; Vianna *et al*., 2019). Haemolytic anemia has been reported in intoxications with aluminium (Aggarwal *et al*., 1999; Soltaninejad *et al*., 2011; Arefi *et al*., 2013; Malakar *et al*., 2019), lead (Warang *et al*., 2017) and mercury (Ribarov *et al*., 1983; Maheswaram *et al*., 2008; Yildirim *et al*., 2012; Ekawanti *et al*., 2015; Weinhouse *et al*., 2017). The anemia is caused when the metallic ions induce membrane damage which predisposes the erythrocytes to intravascular hemolysis or removal from the bloodstream by phagocytes in the reticuloendothelial organs. The basic mechanism of the toxic injury by metallic ions is oxidative stress due to production of reactive oxygen species and free radicals promoted by Fenton reaction (Jaishankar *et al*., 2014). Membrane-bound enzymes which mediate osmoregulation and osmotic stabilization of erythrocytes are altered by these injurious processes (Ribarov *et al*., 1983; Wani *et al*., 2015; Igbokwe *et al*., 2019) and plasma membrane function may be evaluated by erythrocyte osmotic fragility (EOF) technique (Pagano and Faggio, 2015; Farag and Alagawany, 2018).

Osmoregulation in erythrocytes is maintained by transmembrane ion transporters so that osmotic stabilization is achieved by ion leaks into and out of erythrocytes, to provide osmotic equilibrium and avoid osmotic gradient with water flow across the plasma membrane beyond the capacity of regulatory volume adjustments (Armstrong, 2003). Severe toxic injury to erythrocytes interferes with transmembrane ion transporters and inhibits Na^+^-K^+^pump causing intracellular influx of ions with water leading to erythrocyte swelling or oncosis (Vossenkamper and Warnes, 2019) due to necrotic volume increase (NVI) from defect of membrane semi-permeability (Barros *et al*., 2001). Erythrocyte oncosis (erythroncosis) with hydration would reduce the capacity of erythrocytes to withstand hyposmotic stress, thereby increasing EOF (Igbokwe, 2018). In contrast, mild to moderate toxic injury leads to eryptosis where erythrocytes shrink by losing ions and water (Bissinger *et al*., 2019) in eryptotic (apoptotic) volume decrease (EVD) with dehydration (Bortner and Cidlowski, 2002). The eryptotic erythrocytes transform morphologically to echinocytes (Chukhlovin, 1996) and have reduced susceptibility to osmotic fragility (Igbokwe, 2018; Igbokwe *et al*., 2019), with high resistance to osmotic loading (Mindukshev *et al*., 2007). However, spheroechinocytes may have decreased deformability with increase in EOF (Igbokwe, 2018) due to the switch from eryptosis to necrosis and reversion to NVI (Barros *et al*., 2001).

Toxicities involving aluminium and lead ions were reported to increase EOF (Igbokwe, 2018). There was hemolysis of erythrocytes exposed to mercuric ions in isotonic (154 mMol) or slightly hypotonic (150 mMol) saline (Kerek *et al*., 2018) and slightly hypotonic (300 mosmol/L) glucose or sucrose (Igbokwe, 2016; Igbokwe *et al*., 2018), as well as erythrocytes exposed to lead ions in normal saline (Mrugesh *et al*., 2011).The increased fragility reflects the induction of erythroncosis and erythronecrosis (Barros *et al*., 2001; Vossenkamper and Warnes, 2019). On the other hand, decreased EOF was also reported in toxicities of erythrocytes with aluminium and mercuric ions (Igbokwe, 2018). Eryptosis has been reported in erythrocytes exposed to aluminium, lead and mercuric ions (Repsold and Joubert, 2018), which could elicit osmotic resistance by decreasing erythrocyte volume (Bortner and Cidlowski, 2002) and inhibiting water channels to reduce water flux (Igbokwe *et al*., 2018, 2019). Therefore, we hypothesized that the *in vitro* response of goat erythrocytes to cytotoxic injury could cause decreased or increased EOF because of EVD or NVI, respectively (Barros *et al*., 2001; Bortner and Cidlowski, 2002) and subsequently proposed an *in vitro* model of EOF for the assessment of the effect on plasma membrane stability due to eryptosis or erythroncotic necrosis caused by metallotoxic compounds (Figure 1). In this study, we carried out an investigation into the heterogeneous response of goat erythrocytes to osmotic fragility induced during *in vitro* exposure to aluminium chloride, lead acetate or mercuric chloride at low concentration, in order to gain an insight, by means of EOF, into the metallotoxic damage associated with inducible osmotic resistance in addition to elevated osmotic fragility due to erythrocyte death.

## Materials and methods

### *In vitro* cytotoxicity model

The experiment used erythrocytes from goat blood to assess the *in vitro* cytotoxicity of metallic salt solutions (MSS) at low exposure concentrations. Toxic erythrocyte damage by eryptotic or erythroncotic alterations was determined at low and high hyposmolarity of sucrose media. Goat erythrocytes were used in this model because the EOF characteristics in sucrose media were previously described (Igbokwe, 2016; Igbokwe and Igbokwe, 2016) and were presumed to be hypothetically suitable for the investigation of cytotoxic erythrocyte injury that induced EVD and NVI (Fig. 1). Empirical assessment of erythrocyte death modality was based on the outcomes of eryptotic stabilization and erythroncotic destabilization of erythrocyte membranes during EOF (Igbokwe and Igbokwe, 2015; Igbokwe, 2018; Igbokwe *et al*., 2019).

### Source of goat blood

Apparently healthy non-pregnant and non-lactating female Sahel goats, aged about 2.5 years by dentition and weighing 22-25 kg, were selected for this study from the university animal farm. They were kept under semi-intensive management in the farm as earlier described (Igbokwe and Igbokwe, 2015). Blood sample (5ml) was collected from the external jugular vein of each goat in the morning, anticoagulated in heparinised (heparin, 1.0 mg/mL) plastic tubes (Silver Diagnostics, Lagos, Nigeria) and transported on ice to the laboratory where it was analysed within 2 hours. The packed cell volume, erythrocyte count and mean corpuscular volume of each goat were determined and reported as normal parameters previously (Igbokwe and Igbokwe, 2015).

### Preparation of reagents

Isosmotic (isotonic) sucrose (308mosmol/L) stock solution was prepared as described previously (Igbokwe and Igbokwe, 2016) using sucrose (BDH; Poole, England) with a molar mass of 342.3g/mol. The stock solution was diluted to hyposmotic concentrations of 100 and 250 mosmol/L using the procedure that was previously described (Igbokwe and Igbokwe, 2015, 2016). Aluminium chloride (BDH; Poole, England), lead acetate trihydrate (BDH; Poole, England) and mercuric chloride (BDH; Poole, England) have molecular weights of 133.34g, 379.33g and 271.52g, respectively. The metallic salts were used to prepare stock salt solutions with osmotic concentration of 308 mosmol/L (Igbokwe, 2016). Briefly, the procedure for preparing solutions of a specific concentration involved using the molecular weight or molar mass to prepare a molar solution; and thereafter, the osmotic concentration was derived by multiplying the molar concentration with the dissociation factor which was the number of ions in a molecule with presumed complete compound dissociation at high dilution and ambient temperature. The calculated volumes of aliquots were obtained with the formula of equivalence (C_a_*V_a_ = C_b_*V_b_) of the products of concentrations (C) and volumes (V) to achieve appropriate dilutions as earlier reported (Igbokwe and Igbokwe, 2015, 2016).

### Determination of effect of metallic salt solutions on erythrocyte osmotic fragility

The sucrose-based erythrocyte osmotic fragility (EOF) technique was previously described (Igbokwe and Igbokwe, 2015, 2016). This study adopted an abridged EOF with hyposmotic media at 100 and 250 mosmol/L as high (HH) and low (LH) hyposmolarity, respectively. Each metallic salt solution (MSS) was tested with blood samples from 10 goats. The EOF for each blood sample was set up in six text tubes (TT 1-6) with admixtures of hyposmolar media, MSS and aliquot of blood sample as summarized in Table 1. In the set-up, TT1-2 and TT3-4 tested the EOF of the blood at 100 and 250 mosmol/L with aluminium chloride (75 μmol/L), or lead acetate (100 μmol/L) solution at an added osmotic concentration of 0.3 mosmol/L or mercuric chloride solution (133 μmol/L) at an added concentration of 0.4 mosmol/L in TT2 and TT4. The TT5 and TT6 contained isosmotic sucrose medium and deionized distilled water, respectively. The test tube contents were mixed after each test tube received 5 μL of the blood and allowed to incubate at room temperature (35-38 °C) for 30 min. The contents of the tubes were centrifuged at 3000 × g for 15 min; the supernatant of the hemolysate in each tube was harvested with a suction pipette into a cuvette, and the colour of hemoglobin was estimated as absorbance units with a spectrophotometer (ALL PRO; Shibei, Qingdao, China) set at 540 nm, using the supernatants of the tubes containing isosmotic solution (TT5) and deionised distilled water (TT6) as blank (0%) and complete (100%) hemolysis, respectively (Igbokwe and Igbokwe, 2016). The estimate of the EOF, as percentage hemolysis, at each haemolytic endpoint was calculated with a formula (Igbokwe and Igbokwe, 2015):

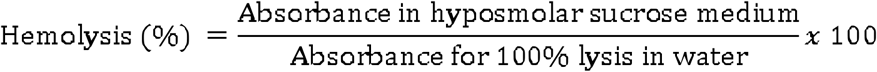

**Table 1.** Experimental testing for the effect of metallic ions on the osmotic stability of erythrocytes of goats in test tubes containing (+) sucrose media, deionized distilled water, metallic salt solution and blood

| Test Tube | Sucrose media (mosmol/L) |  |  | Deionized distilled water<br>(5 mL) | *Metallic salt solution<br>(5/7 µL) | Blood<br>(5 µL) |
| --- | --- | --- | --- | --- | --- | --- |
|  | 100 | 250 | 308 |  |  |  |
|  | (5 mL) | (5 mL) | (5mL) |  |  |  |
| 1 | + | - | - | - | - | + |
| 2 | + | - | - | - | + | + |
| 3 | - | + | - | - | - | + |
| 4 | - | + | - | - | + | + |
| 5 | - | - | + | - | - | + |
| 6 | - | - | - | + | - | + |

\*Metallic salts solutions (308 mosmol/L) was 5 µL for aluminium chloride and lead acetate (0.3 mosmol/L, 75 and 100 µmol/L) and 7 µL for mercuric chloride (0.4 mosmol/L, 133 µmol/L).

### Calculations relating to erythrocyte stabilization and destabilization

Osmotic stabilization (%) or destabilization (%) of erythrocytes of each goat blood induced by MSS was calculated as:

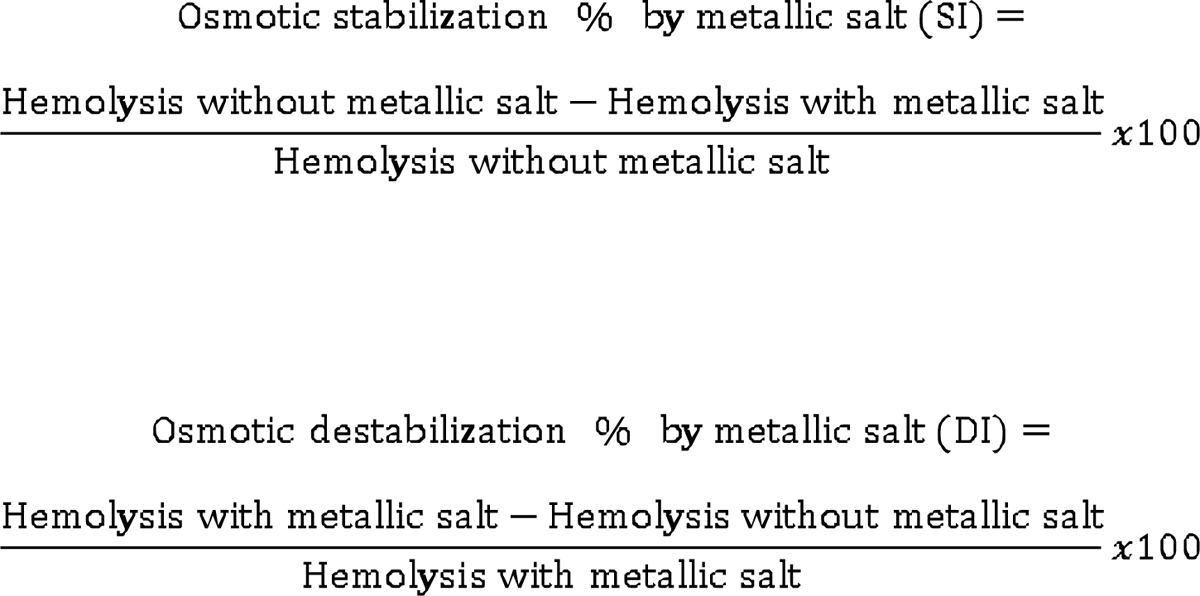

The means of SI or DI at both HH and LH media and the number of animals involved were used to calculate the pooled mean stabilization (Sm) or destabilization (Dm) by each MSS:

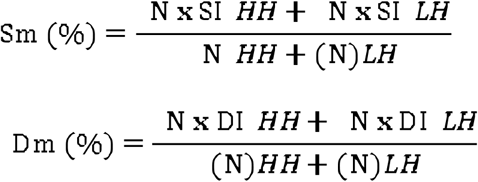

The inducible osmotic resistance (IOR) index of MSS in HH and LH media was calculated as the ratio of Sm to Dm (Sm/Dm).

The probability (P) of occurrence of stabilization (S) or destabilization (D) alone or together induced by MSS was calculated as:

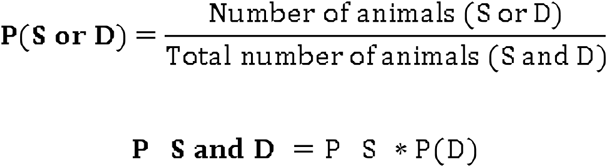

### Statistical analysis

The data were presented as proportions (ratios) or percentages. Summarized data were means with standard deviations, and means were compared with Student’s t-test or analysis of variance with Tukey posthoc test using computer software (GraphPad Instat).

## Results

### Effect of aluminium chloride

The estimates of osmotic hemolysis of erythrocytes in hyposmolar sucrose media containing aluminium chloride are summarized in Table 2. The mean haemolytic estimate induced by the MSS was increased (p < 0.05) from normal value in LH media, but no variation from normal value occurred in HH media. The MSS induced stabilization (□3.5%) in HH media and destabilization (□88.1%) in LH media with the 10 blood samples. The stabilization was much lower (p < 0.05) than the destabilization in HH and LH media.

**Table 2:** Osmotic lysis of goat erythrocytes in hyposmotic sucrose media containing aluminium chloride

| Concentration<br>of sucrose in<br>media<br>(mosmol/L) | Hemolysis (%) in media with<br>aluminium chloride (mosmol/L) |  | Osmotic<br>stabilization<br>(%) | Osmotic<br>destabilization<br>(%) |
| --- | --- | --- | --- | --- |
|  | 0.0 | 0.3 |  |  |
| 100 | 94.4±3.1<sup>a</sup> | 91.2±6.5<sup>a</sup> | 3.5±4.3 (n=10) | 0.0 (n=0) |
| 250 | 8.5±0.7<sup>a</sup> | 71.1±8.2<sup>b</sup> | 0.0 (n=0) | 88.0±0.8 (n=10) |

<sup>a,b</sup>Means±standard deviations with different superscripts are significantly (p<0.05) different

### Effect of lead acetate

The hemolytic estimates induced by lead acetate in HH and LH media are presented in Table 3. The MSS induced both stabilization (5.5%) and destabilization (7.7%) in HH media, but induced only destabilization (89.8%) in LH media. The hemolytic estimate was higher (p < 0.05) than normal value in LH media, but no variation from normal value occurred in HH media. The stabilization (5.5%) was lower (p < 0.05) than the destabilization (89.8%).

**Table 3:** Osmotic lysis of goat erythrocytes in hyposmotic sucrose media containing lead acetate

| Concentration<br>of sucrose<br>in media<br>(mosmol/L) | Hemolysis (%) in media with lead<br>acetate (mosmol/L) |  | Osmotic<br>stabilization<br>(%) | Osmotic<br>destabilization<br>(%) |
| --- | --- | --- | --- | --- |
|  | 0 | 0.3 |  |  |
| 100 | 92.6±5.8<sup>a</sup> | 89.5±5.9<sup>a</sup> | 5.5±3.2 (n=8) | 7.7±0.8 (n=2) |
| 250 | 8.5±0.7<sup>a</sup> | 84.3±8.1<sup>b</sup> | 0.0 (n=0) | 89.8±1.5 (n=10) |

<sup>a,b</sup>Means±standard deviations with different superscripts are significantly (p<0.05) different

### Effect of mercuric chloride

In Table 4, the exposure of erythrocytes to mercuric chloride in HH media caused a significant (p < 0.05) reduction of mean hemolytic estimate from normal value, but no variation from normal value occurred in LH media. In HH media, there was stabilization (95.1%) with all blood samples without any occurrence of destabilization. In LH media, both stabilization (12.2%) and destabilization (21.0%) occurred.

**Table 4:** Osmotic stabilization of Sahel goat erythrocytes in sucrose media containing mercuric chloride

| Concentration<br>of sucrose<br>in media<br>(mosmol/L) | Hemolysis (%) in media with<br>mercuric chloride<br>(mosmol) |  | Osmotic<br>stabilization<br>(%) | Osmotic<br>destabilization<br>(%) |
| --- | --- | --- | --- | --- |
|  | 0.0 | 0.4 |  |  |
| 100 | 94.4±2.8<sup>a</sup> | 5.19±0.9<sup>b</sup> | 95.1±1.8 (n=10) | 0.0 (n=0) |
| 250 | 2.08±0.9<sup>a</sup> | 1.92±0.6<sup>a</sup> | 12.2±12.3 (n=7) | 21.0±9.1 (n=3) |

<sup>a,b</sup>Means±standard deviations with different superscripts are significantly (p<0.05) different

### Inducible osmotic resistance (IOR) by the metallic salt solutions (MSS) in hyposmolar sucrose media

The IOR and probability of erythrocyte stabilization or destabilization induced by each MSS are summarized in Table 5. The Sm was 61.0% with mercuric chloride, and 5.5% and 3.5% when lead acetate and aluminium chloride were tested, respectively. The IOR values were 2.90, 0.07 and 0.04 for the mercuric, lead and aluminium compounds, respectively. The Dm by aluminium chloride was 88.0%, but it was 76.1% and 21.1% when lead acetate and mercuric chloride were tested, respectively. The probability of mercuric chloride inducing stabilizing effect in goat erythrocytes was 0.85, but it was 0.40 and 0.50 for lead acetate and aluminium chloride, respectively. The probability of a destabilization was high with aluminium chloride (0.50) and lead acetate (0.60), but it was low with mercuric chloride (0.15). The probability of concurrence of both effects was 0.13, 0.25 and 0.26 with mercuric chloride, aluminium chloride and lead acetate, respectively.

**Table 5:** Erythrocyte stabilization or destabilization by aluminium chloride, lead acetate or mercuric chloride

| Variable | Aluminium chloride | Lead acetate | Mercuric chloride |
| --- | --- | --- | --- |
| Mean stabilization, Sm, % | 3.5 | 5.5 | 61.0 |
| Mean destabilization, Dm, % | 88.0 | 76.1 | 21.1 |
| Inducible Osmotic resistance, IOR | 0.04 | 0.07 | 2.90 |
| Probability (P) of occurrence (ratio): |  |  |  |
| Stabilization, P(S) | 0.50 | 0.40 | 0.85 |
| Destabilization, P(D) | 0.50 | 0.60 | 0.15 |
| Both S & D, P (S*D) | 0.25 | 0.24 | 0.13 |

**Table 6:** Comparison of erythrocyte stabilization and destabilization in high and low hyposmolar sucrose media

| Parameters for all<br>metallic salt<br>solutions | Hyposmolar sucrose media |  |
| --- | --- | --- |
|  | High, 100 mosmol/L | Low, 250 mosmol/L |
| Stabilization: |  |  |
| Pooled mean (Sp), % | 36.8 (3.5-95.1) | 7.7 (0.0-7.7) |
|  | [n =28] | [n = 2] |
| Increase in Sp, % | 79.1 | -- |
| Probability (S), ratio | 0.93 (28/30) | 0.07 (2/30) |
| Increased probability (S), ratio | 13.3 (0.93/0.07) | -- |
| Destabilization: |  |  |
| Pooled mean (Dp), % | 12.2 (0.0-12.2) | 80.0 (21.0-88.0) |
|  | [n =7] | [n = 23] |
| Increase in Dp, % | -- | 84.7 |
| Probability (D), ratio | 0.23 (7/30) | 0.77 (23/30) |
| Increased probability (D), ratio | -- | 3.3 (0.77/0.23) |

### Comparison of erythrocyte osmotic lysis induced by metallic salt solutions (MSS) in high and low hyposmolar sucrose media

Erythrocyte stabilization or destabilization induced by MSS in high and low hyposmolar media are presented in Table 7. Stabilization of erythrocytes induced by MSS was 79.1% higher in HH than LH media. The stabilizing effect was observed when 28 out 30 goat blood samples were tested. The probability of stabilization as an outcome of the test was 0.93 in HH media against 0.07 in LH media. Stabilization had 13.3 folds of the chance of occurrence than destabilization in HH media. Only 2 samples (20%) manifested a stabilization of 7.7% in HH media that was induced by only lead acetate. Destabilization of erythrocytes, on the other hand, was induced by MSS with 23 out of 30 goat blood samples and the level was 84.7% higher in LH than HH media. The destabilization occurred with a higher probability of 0.77 than the probability of 0.23 for stabilization in LH media. Destabilization had 3.3 folds of chance to occur instead of stabilization in LH media. Only mercuric chloride had the capacity to destabilize erythrocytes in LH media with 70% of blood samples tested.

## Discussion

This study demonstrated the application of abridged sucrose-based EOF in xenobiotic cytotoxicity testing. The theoretical model (Figure 1) was validated by the observation of increased or decreased EOF induced by the MSS. Our data demonstrated strong erythrocyte stabilization and destabilization in HH and LH media, respectively. The phenotypic reasons for the construct of this EOF model were based on EOF characteristics of goat erythrocytes (Igbokwe and Igbokwe, 2015, 2016), which was a sigmoidal curve of dependence of hemolytic estimates on the hyposmolarity of sucrose media (Igbokwe and Igbokwe, 2015), as was similarly affirmed in a subsequent report (Singh *et al*., 2019). The sucrose media stabilized goat erythrocytes from median to maximal hyposmolarity (Igbokwe and Igbokwe, 2015). The variable non-hemolytic hyposmolar media concentrations of sucrose were 240-300 mosmol/L (Igbokwe and Igbokwe, 2016). The LH media concentration here was 250 mosmol/L, so that the counteracting destabilizing effect would be demonstrable during xenobiotic exposure. The hemolysis in LH media was ≤ 8.5% for the control and depicted anticipated stabilizing effect, but this effect was non-existent at hyposmolar concentrations (<120 mosmol/L) causing >90% hemolysis (Igbokwe and Igbokwe, 2015, 2016). The hyposmolar concentration of HH media at100mosmol/L elicited hemolytic estimates of ≥ 92.6% for controls showing maximum erythrocyte destabilization, so that stabilizing effect could be demonstrated during erythrocyte cytotoxicity.

For each MSS, the induced change in EOF was both an increase and a decrease depending on the media hyposmolarity and varied with the erythrocytes of individual goats. The metallic toxicants are pro-oxidants causing oxidative injury to erythrocytes (Jaishankar *et al*., 2014). The variability of hemolytic outcomes would depend on the injurious stimulus derived from the balance of generated oxidant load and the vigour of antioxidant systems of the erythrocytes. Other factors affecting the goat-dependent EOF outcomes might include ion contents of erythrocytes, number and functionality of transmembrane ion transporters, erythrocyte volume and metabolic state, and membrane surface-to-volume ratio and composition. These physiological factors affect how the membrane responds to injury and functions as EVD or NVI is induced. The hydration status of the erythrocytes is influenced by the intracellular ion content (Gallagher, 2017), and decreased or increased erythrocyte hydration causes decreased or increased EOF, respectively (Cueff *et al*., 2010). The erythrocytes with membranes that are diffusible to metallotoxic ions are expected to have enhanced metabolic derangement, followed by drastic energy depletion, and less efficient antioxidant systems. Sahel goats have wide variability in erythrocyte glutathione contents with varied capacity for glutathione-based antioxidant defence (Igbokwe *et al*., 1998). The erythrocyte death modality, factored as erythrocyte fragility influencer, would be dependent on disposable energy and the capacity of glutathione regeneration in erythrocytes (Kurata *et al*., 1994; Orrenius *et al*., 2011), and these factors determined whether the injury of the goat erythrocytes responded with increased or decreased EOF. Therefore, the dual EOF outcomes (stabilization and destabilization) of goat erythrocytes probably depended on physiological characteristics of erythrocytes and their ability to cope with physiological risks associated with toxic injury, so that attenuated risks elicited EVD and osmotic stabilization while uncontrolled risks led to NVI and osmotic destabilization.

Mercuric chloride induced high osmotic resistance, whereas aluminium chloride and lead acetate were potent inducers of osmotic fragility based on IOR. The IOR of ≥1.0 indicated strong osmotic resistance, while strong hemolytic potential was indicated by a value near zero. Apparently, each MSS had the capacity to induce both osmotic resistance and fragility under differentiated hyposmolar environments. The dual EOF outcomes induced by the MSS suggested that cytotoxic erythrocyte death programme could be detected by this simple EOF technique. The toxic ions might have interacted with death receptors on surfaces of erythrocyte membranes or entered the erythrocytes through ion channels by active transport (Bridges and Zalups, 2017), passive diffusion (Simons, 1986) or both transport modes (Exley and Mold, 2015). Extensive intracellular traffic of toxic ions was expected to cause severe damage to metabolic processes that would undermine cellular energy output and antioxidant capacity, and this could predispose intoxicated erythrocytes to severe homeostatic disruption leading to erythrocyte necrosis (hemolysis) as observed with toxic actions where EOF increased. Mild cytotoxic injury would arise when intracellular content of the toxicant was low; with the consequence that metabolic energy depletion was minimized and antioxidant activity was not overwhelmed, so that residual cellular energy could be utilized to initiate energy-dependent eryptosis (Orrenius *et al*., 2011). Delayed cellular uptake of mercuric ions in media was reported in erythrocytes, because the ions were bound to membrane proteins without crossing the membrane into the cytoplasm (Zolla *et al*., 1994). The EOF depended on erythrocyte swelling, membrane stretching and holes formation, but the primary swelling kinetic would be influenced by mercuric ion-sensitive water channels on membranes (Pribush *et al*., 2002). The mercuric ions were reported to cross the erythrocyte membranes later into the cytoplasm to cause hemolysis (Zolla *et al*., 1994). Thus, IOR was probably associated with low cytoplasmic distribution of toxic ions with less severe cytotoxicity causing EVD, along with inhibition of primary swelling by reduced activity of water channels on erythrocyte membranes (Bortner and Cidlowski, 2002; Pribush *et al*., 2002; Orrenius *et al*., 2011).

Mercuric chloride induced increased EOF and erythrocyte destabilization in LH media with low probability and moderate intensity, consistent with reports of mercuric ion-induced hemolysis in slightly hyposmolar or isosmolar media (Kerek *et al*., 2018; Igbokwe, 2016; Igbokwe *et al*., 2018). Aluminium chloride and lead acetate had more destabilizing effects in LH than HH media. Erythrocyte destabilization was more likely in LH than HH media probably because of molecular crowding with sucrose that facilitated interaction of toxic ions with the erythrocytes to enhance the toxic actions (Takahashi *et al*., 2020). Where destabilization by the MSS occurred in HH media, it was likely due to subsequent permeabilization of echinocytes after transformation to spheroechinocytes or eryptotic microvesicles (Kerek *et al*., 2018). The reduction in crowding of toxic ions with sucrose in HH media would have greatly reduced toxic activity in cytoplasmic milieu and prevented severe cellular injury with hemolytic outcomes, but erythrocyte destabilization in HH could be due to postapoptotic necrosis (Orrenius *et al*., 2011).

The EOF model produced increasing hemolysis from low to high hyposmolarity in a monophasic manner considered as monotonic (Singh *et al*., 2019). As the toxic action increased with molecular crowding, the toxic response was expected to be monotonic, but it seemed to deviate from monotonicity since EOF outcomes diverged at both ends of hyposmolarity spectrum. This represented a nonmonotonic response probably influenced by the magnitude of toxic actions in HH and LH media (Conolly and Lutz, 2004). The use of EOF in cytoxicity testing had been recognized (Pagano and Faggio, 2015; Farag and Alagawany, 2018) without positing the dimensions of the dual opposing outcomes that could be generated from the test as revealed from this study. The cytotoxicity of toxicants in erythrocytes ought to include eryptosis and necrosis, with EOF as a preliminary measuring tool for these death modalities, but EOF had not been hitherto considered as relevant to the assessment of erythrocyte death (Cumming *et al*., 2012).

In this study, we identified the exposure level to be close to “no observed adverse effect level” (NOAEL) because no significant differences from control were induced by aluminium chloride or lead acetate in HH media and mercuric ions in LH media. However, the EOF outcomes from individual goats exposed to lead and mercuric ions expressed osmotic resistance and fragility as variants to the adverse effects. The no-observed effect was a nullification of opposing amplitudes in the toxic action which was made obvious by extension of the exposure level to lower levels (Igbokwe, 2016) or further reduction of hyposmolarity (Igbokwe *et al*., 2018). The biphasic or dual outcome with similar quantitative features across the phases was a subtle pointer to a hormetic threshold indicated by the EOF model. Hormesis as a biphasic response characterized by low-dose stimulation and high-dose inhibition extends through NOAEL dose where there is neutrality (Calabrese and Baldwin, 2002, 2003). The EOF, as toxicological endpoint, can indicate stimulation of eryptosis through increased IOR, which is a physiological imperative for osmotic stability; and inhibition connoted decreased IOR with increasing EOF as the usual expectation in adverse effect. Chemical hormesis existed in the toxic actions of aluminium and mercuric ions (Calabrese and Baldwin, 2003). The adaptive hormetic osmotic resistance prevents hemolysis as the erythrocytes circulate in the bloodstream, and within physiological limits may be moderated by antioxidant actions that antagonize toxic effect. Thus, EVD-induced osmotic resistance may be corrected by regulatory volume increase to restore erythrocyte viability, but erythrocytes may also be irreversibly dehydrated by solute loss to the extent that regulatory volume adjustment is no longer feasible, and the erythrocytes become senescent (Gallagher, 2017). Erythrocytes, with induced osmotic resistance, have membrane surface receptors that bind to phagocytes for their removal from the bloodstream and for engagement in intravascular coagulation (Qadri *et al*., 2017). When the osmotic resistance is inhibited, EOF increases with hemoglobin leak through membrane holes, complete membrane rupture (Pribush *et al*., 2002) or engulfment of oncotic erythrocytes with exposed receptors for their capture by phagocytes (Lecoeur *et al*., 2001).

## Conclusion

The *in vitro* cytotoxicity model (using EOF) evaluated the toxic actions of aluminium chloride, lead acetate and mercuric chloride at low concentrations (around NOAEL) which affected osmotic membrane stability in varied hyposmolar environments, and it showed that the toxic actions induced dual outcomes of osmotic resistance (stabilization) and fragility (destabilization) concurrently, such that the monotonic EOF could be deviated by the toxic effect. The likelihood of osmotic stabilization or destabilization was increased by high or low media hyposmolarity, respectively. The erythrocytes of individual goats responded to toxic actions with dual EOF outcomes where no observed adverse effect was encountered in the group. Thus, EOF could be an applicable test in toxic conditions causing EVD from eryptosis or NVI from erythronecrosis, and the test model might be used to gauge dual or biphasic, nonmonotonic and hormetic outcomes in cytotoxicity of erythrocytes.

## Acknowledgements

Financial assistance to Nanacha Afifi Igbokwe was granted by the Council of the University of Maiduguri through a study fellowship for postgraduate training. Technical assistance was provided by the research laboratory of the Department of Veterinary Pathology, University of Maiduguri, Maiduguri, Nigeria.

## Ethical Compliance

Approval for the research (Ref: PGA/12/07640) was granted by the Board of School of Postgraduate Studies, University of Maiduguri, Maiduguri, Nigeria, after its ethical committee affirmed that the research complied with institutional, national and international guidelines on the use of animals and animal resources for research.

## Legend to Figure 1

**Figure 1:** The theoretical model of how cytotoxic injury causes anemia by eryptosis with eryptotic volume decrease (EVD) or erythroncotic necrosis with necrotic volume increase (NVI) [A]; and the erythrocyte osmotic fragility model used in testing erythrocyte cytotoxicity [B]

